# Genomic Characterization of a Locally Transmitted *Leishmania mexicana* Isolate from Texas

**DOI:** 10.1101/2025.07.31.668039

**Authors:** Juan David Ramírez, Luz H. Patiño, Binita Nepal, Sarah M. Gunter, Eva H. Clark, Dawn M. Wetzel

## Abstract

Cutaneous leishmaniasis (CL) is caused by *Leishmania* species transmitted by sand flies. Although considered a neglected tropical disease, growing evidence indicates that local transmission can occur in subtropical, higher-resource regions where sand fly vectors are present, including the southern United States. Here, we report the first whole-genome sequence of *Leishmania mexicana* from an autochthonous U.S. case—a 3-year-old boy from Ellis County, Texas, with no travel history. Genomic DNA was extracted from a skin biopsy specimen and sequenced using Illumina technology. Phylogenomic analysis based on nuclear and mitochondrial SNPs confirmed the isolate as *L. mexicana*, clustering with other members of the *L. mexicana* complex. The parasite exhibited a predominantly disomic karyotype, with chromosome 30 displaying trisomy. We identified 172 genes with significant copy number variations, including genes involved in ubiquitination, nucleic acid metabolism, and protein translation. Additionally, 53,964 SNPs were detected, over 22,000 of which were predicted to have moderate or high functional impact, affecting genes linked to host-pathogen interactions, metabolic pathways, and signal transduction. This study provides the first genomic characterization of a locally acquired *L. mexicana* strain in the U.S. and underscores the value of molecular surveillance and increased clinical awareness of leishmaniasis in subtropical regions where competent vectors are present.

## IMPORTANCE

Leishmaniasis is traditionally associated with tropical and low-resource settings, yet increasing reports of autochthonous cases in the southern United States highlight the need to recognize its presence in non-endemic regions. This study provides the first whole-genome sequence of *Leishmania mexicana* from a locally acquired case in the U.S., offering valuable insights into the parasite’s genetic makeup and potential adaptations. By identifying chromosomal and gene-level variations, including mutations in genes related to host interaction and metabolism, our findings contribute to a growing body of knowledge on *Leishmania* evolution and biology in new geographic contexts. These data support the use of genomic tools to enhance surveillance, inform public health strategies, and improve clinical recognition of leishmaniasis in regions where sand fly vectors are established but often overlooked.

## OBSERVATION

Leishmaniasis is a vector-borne disease caused by protozoan parasites of the genus *Leishmania*. This pathogen has a global distribution with endemic transmission found in tropical and subtropical regions (1). In the Americas, there has been endemic transmission noted in 21 countries predominately in Central and South America (2, 3). Sporadic autochthonous cases have been reported in the southern US, with the major disease burden in Texas. Environmental changes caused by climate change will likely continue to alter the epidemiology of this disease and increase the burden in the US, particularly in Texas and Oklahoma (4, 5). Sand fly vectors, long present in the region, appear to be expanding their geographic range, and *Leishmania mexicana* has been identified in local animal reservoirs, supporting ongoing local transmission in Central and Northern Texas (5, 6). Most U.S. clinicians are unaware that cutaneous leishmaniasis (CL) can be caused by local *Leishmania* species, and consequently only include CL in their differential diagnosis of characteristic skin lesions in the context of foreign travel or migration (6-8).

Previously, we published a case series of three pediatric patients with cutaneous leishmaniasis (CL) and no history of travel that were evaluated at the Pediatric Infectious Diseases Clinic at the University of Texas Southwestern Medical Center in Dallas within a 6-month period (9). All three children resided in North Texas, in areas where local sand fly vectors are known to be present. Lesions were non-healing and ulcerative, and diagnosis was delayed in each case due to initial misclassification as bacterial or inflammatory skin disease. Histopathology and PCR of the internal transcribed sequence 2 region of ribosomal DNA (ITS2) confirmed *L. mexicana* infection in each instance.

We now describe the first whole-genome sequence of *Leishmania mexicana* from a locally acquired human case in the U.S., obtained from one of the previously reported patients with available tissue. The patient was a 3-year-old boy from Ellis County, Texas, who developed a nodular lesion on his arm that persisted and worsened over a 5-month period before a diagnosis was made. The lesion initially progressed after corticosteroid treatment. Histopathology revealed numerous intracellular amastigotes, and molecular testing at the Centers for Disease Control and Prevention (CDC) confirmed *L. mexicana* infection. The patient was treated with fluconazole for 10 weeks, resulting in complete resolution of the lesion.

We extracted genomic DNA from a portion of the skin biopsy specimen and performed whole-genome sequencing using the Illumina NovaSeq 6000 platform (single-end, 150 bp reads). We prepared libraries using the Nextera DNA Flex kit according to the manufacturer’s protocol. We mapped the resulting reads to reference genomes and evaluated the chromosome and Gene CNV, and genetic variation as detailed in **Supplementary File 1**.

Two alignments were used to perform the phylogenomic analyses: one based on SNPs from the nuclear genome and the other on SNPs from the mitochondrial genome. The results show a close relationship between the nuclear and maxicircle SNPs of the newly sequenced sample and the reference genomes of *Leishmania mexicana* complex. Overall, three well-supported clusters were identified: Cluster 1 (highlighted in light red) includes the genomes of *L. infantum* and *L. donovani*; Cluster 2 (light green) includes species of the subgenus *Viannia* (*L. panamensis, L. peruviana, L. braziliensis, L. guyanensis, L shawi, L. naiffi and L. lainsoni)* and Cluster 3 (light blue) groups *L. mexicana, L. amazonensis, L. pifanoi*, and the genome analyzed in this study **(Fig. 1A and 1B)**.

**Fig 1.**
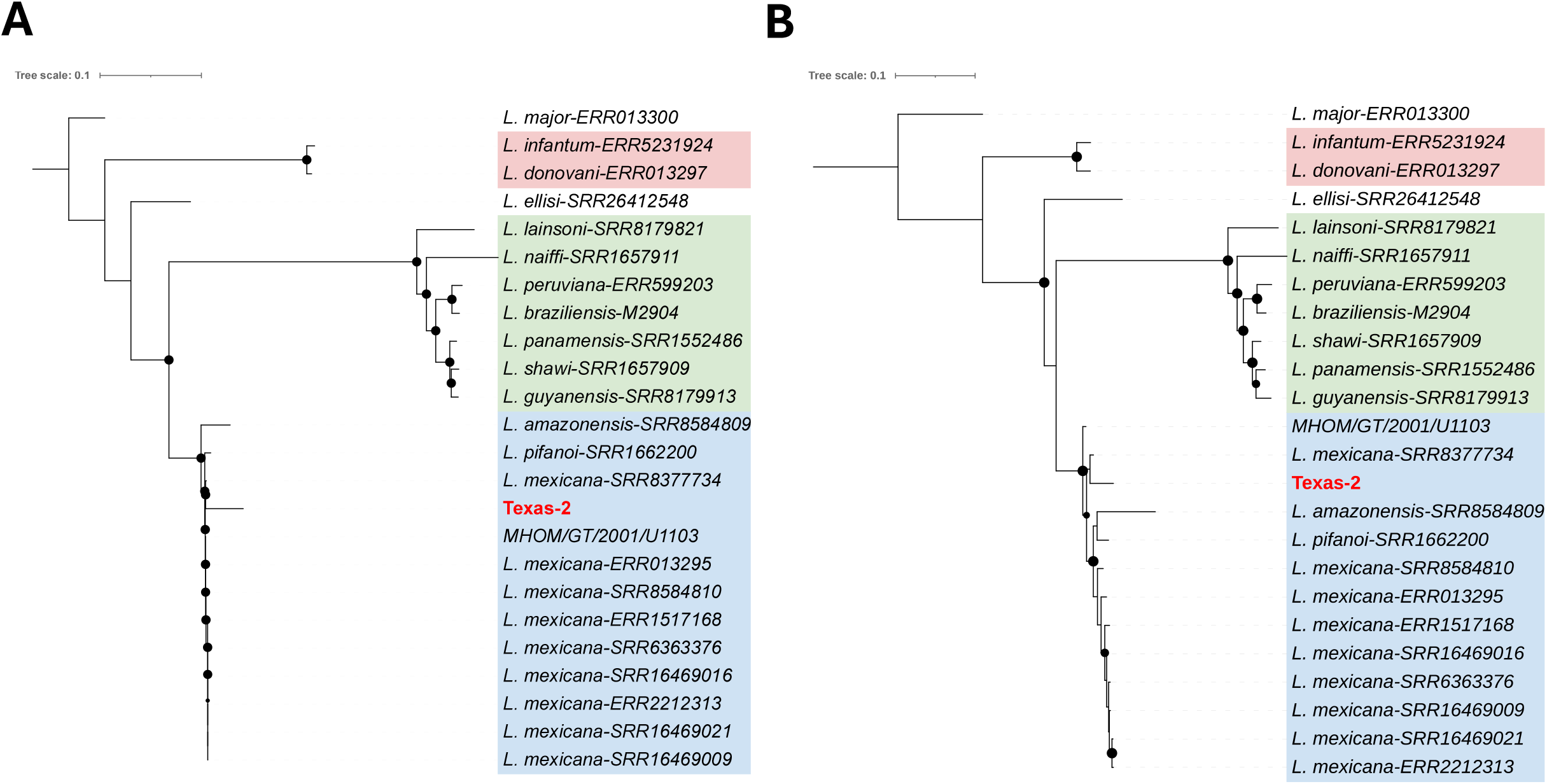
Phylogenetic relationships among *Leishmania* species based on nuclear and mitochondrial SNPs. This figure shows a phylogenomic analysis using SNP data from both nuclear (A) and mitochondrial (B) genomes. It includes species from the *L. mexicana* complex (light blue), the subgenus *Viannia* (light green), the subgenus *Leishmania* (light red), and the sample analyzed in this study (Texas-2). *L. mexicana* MHOM/GT/2001/U1103 was used as the reference strain and *L. major* as the outgroup. Black dots mark well-supported nodes with bootstrap values ≥ 95.

We analyzed and compared the genome’s chromosome copy numbers. The karyotype remained stable, with most chromosomes displaying an S value between 1.5 and 2.0, consistent with a disomic state. The only exception was chromosome 30, which showed a trisomic profile.

To identify genes with copy number variations (CNVs) and assess their occurrence in the genome analyzed, we calculated CNVs for each gene using a z-score cutoff > 2 and an adjusted p-value < 0.05 (10). The results obtained showed 172 genes with CNVs **(Table S1)**. Interestingly, the genes with the highest copy number variations (CNVs) and known functions were *polyubiquitin* and *P1/S1 nuclease*, with z-scores of 31.716 and 21.297, respectively. These were followed by genes involved in the translation machinery, including *TMA7*, ubiquitin-fusion protein, alpha and beta tubulin, and *heat shock protein 83-1* (**Table 1**). Gene Ontology enrichment analysis of CNV-associated genes revealed enrichment in biological processes such as chromatin organization and remodeling, amino acid transmembrane transport, autophagy, biosynthetic and metabolic pathways, and cellular homeostasis. While the direct clinical implications of these CNVs require further investigation, several of the affected genes particularly those involved in stress response (e.g., *HSP83-1*) and chromatin remodeling may be relevant to parasite adaptability, virulence, or potential drug resistance mechanisms (11-13).

**Table 1.** List of genes that presented the highest copy number variation (CNV) in the genome analyzed.

| Gene ID | Start gene | End gene | Gene Annotation | z_score |
| --- | --- | --- | --- | --- |
| LmxM.36.3530partial | 1352798 | 1353040 | polyubiquitin, putative | 31,716 |
| LmxM.29.1510 | 508877 | 509827 | p1/s1 nuclease | 21,297 |
| LmxM.09.0891 | 336699 | 337085 | polyubiquitin, putative | 9,931 |
| LmxM.29.2760 | 1005005 | 1005154 | Translation machinery associated TMA7, putative | 9,931 |
| LmxM.32.0794 | 284398 | 285879 | beta tubulin | 7,090 |
| LmxM.21.1860 | 730734 | 732065 | beta tubulin | 6,142 |
| LmxM.32.0792 | 281233 | 282564 | beta tubulin | 6,142 |
| LmxM.13.0390 | 93414 | 95060 | alpha tubulin | 5,195 |
| LmxM.30.1900 | 908400 | 908786 | ubiquitin-fusion protein | 5,195 |
| LmxM.08.1230 | 1664002 | 1665246 | beta tubulin (fragment) | 4,248 |
| LmxM.32.0312 | 108760 | 110865 | heat shock protein 83-1 | 4,248 |
| LmxM.32.0314 | 112338 | 114443 | heat shock protein 83-1 | 4,248 |
| LmxM.32.0316 | 115917 | 118022 | heat shock protein 83-1 | 4,248 |

We analyzed the number of SNPs in the genomes studied and compared them to the reference *L. mexicana* genome. A total of 53,964 SNPs were identified. When assessing their potential functional impact, approximately 22,040 SNPs were classified as having either moderate or high functional effects, 21,211 SNPs (96%) were predicted to have a moderate impact, while 826 SNPs (4%) showed a high impact (14, 15). Most of the high impact SNPs were associated with *stop gained* mutations and were found in both genes encoding hypothetical proteins and those with known functions. We highlight those SNPs located in genes encoded transporter proteins (pteridine transporter, ABC1 transporter, amino acid and nucleoside transporter protein), host-pathogen interaction associated proteins (amastin surface protein), kinetoplast-associated protein, or intracellular degradation-associated proteins (ubiquitin conjugating enzyme putative), structural proteins (tubulin, kinesins), as well as in genes associated with the intracellular signal pathway, such as phosphatidylinositol 3 and 5 kinase and in glycolysis and gluconeogenesis as glucose 6 phosphate isomerase **(Table S2)**.

This study presents the first whole-genome sequence of *Leishmania mexicana* from an autochthonous U.S. human CL case. This case emphasizes the importance of including CL in the differential diagnosis of patients with characteristic skin lesions who have resided in U.S. regions where sandflies are present (4). Our genomic analysis of one Texas isolate provides high-resolution insight into the molecular characteristics of this local strain and highlights its phylogenetic relatedness to other members of the *L. mexicana* complex (16, 17). Using nuclear and mitochondrial SNP-based phylogenomics **(Figure 1)**, we confirmed that this genome clusters tightly with *L. mexicana, L. amazonensis*, and *L. pifanoi*, distinct from other major *Leishmania* lineages.

Chromosome-level analysis revealed a predominantly disomic karyotype, with the exception of chromosome 30, which displayed a trisomic profile—consistent with the well-documented chromosomal plasticity observed in *Leishmania* spp (18). Notably, we identified 172 genes with significant copy number variations (CNVs), including amplifications in *polyubiquitin, P1/S1 nuclease*, and components of the translational machinery such as tubulins and *heat shock protein 83-1*. These CNVs may represent adaptive responses to host-induced stress or immune pressures and could have implications for parasite survival, virulence, or drug response, although their precise functional roles remain to be elucidated (19-22). Functional enrichment analysis indicated that CNV-associated genes were involved in pathways related to chromatin remodeling, autophagy, and cellular homeostasis, suggesting selective pressures on genome organization. Additionally, SNP analysis identified over 53,000 variants relative to the reference genome, with approximately 22,000 predicted to have moderate or high functional impacts. Among the high-impact variants were genes implicated in host-pathogen interactions (e.g., *amastin* surface proteins), intracellular protein turnover (e.g., ubiquitin-conjugating enzymes), and essential metabolic pathways such as glycolysis (e.g., *glucose-6-phosphate isomerase*). Mutations affecting transporter proteins and signal transduction pathways may influence parasite fitness, drug susceptibility, or virulence, underscoring the potential clinical relevance of these genomic alterations (23-25).

Taken together, our findings provide a genomic blueprint of a locally acquired *Leishmania mexicana* isolate and underscore the value of molecular characterization for understanding parasite diversity and potential adaptations. In the context of increasing reports of autochthonous leishmaniasis in the U.S., this study highlights the importance of clinical awareness regarding local *Leishmania* transmission and supports the integration of genomic tools into surveillance frameworks for neglected tropical diseases emerging in non-traditional settings.

## DATA AVAILABILITY

The whole genome sequences have been deposited in the ENA-NCBI database under accession number PRJEB94538

## Supplementary Material

Supplementary File 1. Materials and Methods

Table S1. List of genes with CNV (z score cut off >2 and adjusted p-value) in the genome analyzed.

Table S2. List of SNPs with high functional impact (stop gained) in the genome analyzed.

